# Sex-specific exploration accounts for differences in valence learning in male and female mice

**DOI:** 10.1101/2025.08.06.668689

**Authors:** Heike Schuler, Eshaan S. Iyer, Gabrielle Siemonsmeier, Ariel M. Weinbaum, Peter Vitaro, Shiqing Shen, Rosemary C. Bagot

## Abstract

Valence, the quality by which something is perceived as good or bad, appetitive or aversive, is a fundamental building block of emotional experience and a primary driver of adaptive behavior. Pavlovian fear and reward learning paradigms are widely used in preclinical research to probe mechanisms of valence learning but with limited consideration of sex as a biological variable despite known sex differences in neuropsychiatric disorders associated with impaired valence. Here, we compare appetitive-only, aversive-only and mixed-valence cue-outcome Pavlovian conditioning paradigms in male and female mice to dissociate effects of context, valence and salience in a sex-specific manner. Using a data-driven approach to identify behaviors indicative of valence learning in an unbiased manner, we compare task performance between paradigms in male and female mice. We show that while male and female mice acquire appetitive and aversive associations in both single- and mixed-valence paradigms, sex differences emerge in single-valence paradigms. Ultimately, we show that these apparent sex differences in valence learning are driven by non-specific baseline differences in exploratory behavior. Males explore more at baseline, altering their trajectory of cue-reward association acquisition whereas females explore less at baseline, increasing shock facilitated freezing in aversive-only contexts, masking cue discrimination. Overall, our findings illustrate how task design differentially impacts behavioral expression in male and female mice and demonstrate that mixed-valence paradigms afford a more accurate assessment of valence learning in both sexes.

## Introduction

Understanding how emotion is represented in the brain remains a major challenge for modern neuroscience. The complex and subjective nature of human emotional experience challenges efforts to uncover its mechanism. While emotion is complex, valence, the intrinsic attractiveness or aversiveness of a stimulus or state, is fundamental and translates across species (Adolphs et al., 2019; Anderson and Adolphs, 2014; Gündem et al., 2022). Valence guides adaptive behavior: aversive stimuli are avoided, and appetitive stimuli elicit approach. Through associative learning, cues that predict valenced stimuli shape behavior in anticipation of the outcome. Disruptions in valence learning are implicated in a variety of neuropsychiatric disorders (Admon and Pizzagalli, 2015; Chiara, 1999; Lissek et al., 2005). For example, patients with major depressive disorder display negative biases in memory recall (Brittlebank et al., 1993; Douglas and Porter, 2010; Elliott et al., 1997; Maniglio et al., 2014; Noworyta et al., 2021), anxiety disorders are generally defined by maladaptive threat generalization (Lee et al., 2024; Lissek et al., 2014; Vandael et al., 2025), and patients with post-traumatic stress disorder suffer from intrusive memories of traumatic events (Ehlers and Clark, 2000). Notably, sex differences exist in the prevalence, symptom presentation and severity of these disorders (Kuehner, 2003; McLean et al., 2011).

Preclinical Pavlovian conditioning protocols are widely used to study mechanisms of valence learning in rodents. Mice or rats learn to associate either an auditory or visual cue or a context (conditioned stimulus; CS) with an inherently valenced stimulus, such as an electric foot shock (unconditioned stimulus; US). Fear conditioning is widely used as a model of associative valence learning. Reward conditioning protocols are less frequently employed, partially because rodents take substantially longer to form reward than fear associations (Deseyve et al., 2024; Domingues et al., 2025). In both instances, animals learn about a single valence in isolation, yet real world environments are rarely restricted to single valence outcomes. The presence of opposite valence outcomes is known to influence learning, yet this has received little attention of late (Konorski, 1973; Laurent et al., 2022)single valence tasks can conflate cue and context learning and confound valence with salience because valenced outcomes are inherently salient. Despite this, limited research has simultaneously examined appetitive and aversive learning in part due to the scarcity of protocols that dissociate opposing valence processes, especially in mice (Lefner and Moghaddam, 2024; Ray et al., 2022, 2020).

To date, few studies have examined associative learning in mixed valence contexts and even fewer have examined both sexes. The few studies that have considered sex as a biological variable point to potential sex differences in both threat and reward learning (Borkar et al., 2020; Keiser et al., 2017; Lefner et al., 2022). However, findings are inconsistent and limitations in experimental designs preclude a clear understanding of the interaction of sex and valence learning. This is particularly important given prevalent sex differences in valence-related psychopathologies, in which often processing of both positive and negative valences is altered (Kuehner, 2003; McLean et al., 2011). Here, we asked how the presence of more than one valence influences learning and behavioral expression of learning in male and female mice to comprehensive profile sex differences in valence learning. We trained male and female mice in purely appetitive, purely aversive or mixed-valence conditioning tasks and used data-driven behavioral profiling combined with a partial-least squares discriminant analysis to distill learning-relevant behaviors in both sexes. We show that both sexes acquire appetitive and aversive associations, yet task design differentially influences behavioral expression of learning-relevant behaviors in males and females. Examining the evolution of learning across tasks revealed baseline sex differences in exploratory behaviors that mask learned cue discrimination in a context dependent manner. Ultimately, we show that a mixed valence context minimizes sex differences in exploration to more accurately probe valence learning in both sexes.

## Materials and Methods

### Animals

7-week-old male and female C57BL/6J mice (Jackson laboratories, Bar Harbor, ME, USA) were group-housed by sex (n=5/cage) for one week prior to experiments. All mice were maintained at 22-25°C, on a 12h light/dark cycle (lights on at 07:00h), with ad libitum food and water prior to the beginning of the experiments. Animals were food restricted at the start of conditioning and maintained at 85% body weight. All testing occurred throughout the light cycle. All procedures were approved by the Animal Care Committee and conformed to McGill University Comparative Medicine and Animal Resources Centre guidelines.

### Apparatus

Mice were trained in conditioning boxes (15.24 × 13.34 × 12.7 cm; Med Associates Inc., USA) enclosed in sound attenuating chambers outfitted with a programmable audio generator connected to a speaker, a grid floor, an infrared light, a fan for white noise and ventilation, and a food port for liquid reward delivery through a syringe pump. Each food port is equipped with an infrared beam to quantify head entries and a lickometer to quantify licks. Boxes were controlled and data collected by a computer running MED-PC software (Med-Associates Inc., USA).

Sessions were recorded from top view using AniHome (Singh et al., 2019) on Raspberry Pi (Raspbian GNU/Linux 10 (buster)) with infrared camera modules (Raspberry Pi Camera Module 3 NoIR 3) at 800 × 600 pixels resolution and 40 frames per second (FPS). To allow for video-based identification of cue periods, infrared LEDs mounted on top of the operant box were triggered each time a cue was presented. Metal trays underneath the grid floor were filled with 1/8” corncob bedding to prevent reflections of infrared lighting in the recording.

### Classical conditioning protocols

#### Task design considerations

Mice take longer to learn an appetitive association than an aversive association (Deseyve et al., 2024). Repeated exposure to cue-outcome pairings after the association has been acquired can lead to overtraining and habituation in animals, such that a learned behavioral response (e.g., freezing) may no longer be exhibited to the cue (Carroll et al., 2024; Tafreshiha et al., 2021). To balance these considerations, protocols incorporated more appetitive cue-outcome pairings than aversive cue-outcome pairings. Single valence conditioning protocols are designed as a subset of the mixed valence conditioning protocol to support direct comparison between the paradigms and account for differences in training duration between appetitive and aversive single valence conditioning protocols.

#### Conditioning protocols

Animals were transported from the colony to the testing room in their homecage and left to habituate under red light 1h prior to session start. Conditioning was preceded by a single day of context and cue habituation, during which mice were placed in the operant box and exposed to five presentations of each cue without outcomes. Each conditioning session began with a 2 minute habituation period, followed by presentations of auditory cues (CS) in random order separated by a variable inter-trial interval (ITI; average 180 sec). The appetitive CS (CS Reward; CS^R^) was paired with 30 µl of chocolate milk dispensed into the food port. The aversive CS (CS Shock; CS^S^) was paired with a 0.5 mA foot shock delivered through the grid floor for 0.5 seconds. The CS^−^ was not paired with any outcome. Auditory cues were presented for 15 seconds, with outcomes delivered randomly within the final 5 seconds of the cue. Cues were reinforced 80% of the time, as prediction error improves learning (Iordanova et al., 2021). Boxes were cleaned and bedding replaced between animals. One day after completion of conditioning, mice were exposed to a recall session (five presentations of each cue without outcomes) to assess behavioral responses to cues in the absence of outcomes.

##### Mixed valence conditioning

Training began with a mixed-valence training session, followed by two appetitive-only training sessions, repeating this order for 14 daily training sessions. Mixed-valence training sessions consisted of 10 CS^R^, 5 CS^S^ and 5 CS^−^ presentations. Appetitive-only training sessions consisted of 10 CS^R^ and 10 CS^−^. Cue identity was counterbalanced for CS^R^ and CS^S^ between two pure tones (2 kHz, 75 dB or 10 kHz, 63 dB), while CS^−^ was always a clicker (10 Hz, 75 dB).

##### Appetitive single valence conditioning

Appetitive conditioning mirrored appetitive-only training days in the mixed valence conditioning protocol. Training lasted for 14 daily sessions. Each session consisted of 10 CS^R^ and 10 CS^−^. Cue identity was counterbalanced between mice for CS^R^ between two pure tones (2 kHz, 75 dB or 10 kHz, 63 dB), while CS^−^ is a clicker (10 Hz, 75 dB).

##### Aversive single valence conditioning

Aversive conditioning mirrored the aversive components of mixed training days in the mixed valence conditioning protocol. Training lasted for 14 days with training sessions every third day interspersed with two rest days when mice remained in their homecage. Each session consisted of 5 CS^S^ and 5 CS^−^. Cue identity was counterbalanced between mice for CS^S^ between two pure tones (2 kHz, 75 dB or 10 kHz, 63 dB), while CS^−^ is a clicker (10 Hz, 75 dB).

#### Conditioning cohorts

The mixed-valence discovery and validation cohort consisted of 56 (n_*Male*_ = 28, n_*Female*_ = 28) and 18 (n_*Male*_ = 9, n_*Female*_ = 9) mice, respectively. The single-valence cohorts consisted of 16 animals each (n_*Male*_ = 8, n_*Female*_ = 8).

### Behavioral analysis

#### Analysis data

Behavior was evaluated using video-based approaches as well as food-port generated outputs. Whole session video recordings were cropped into 45s clips centered on the cue interval (15s prior to cue onset to 15s post cue offset) for freezing analysis (ezTrack) and detailed behavioral profiling (DeepLabCut and Keypoint-MoSeq). Head entries, head exits (infrared beam) and licks (contact lickometer) were recorded and analyzed as indicators of reward seeking.

#### ezTrack

Freezing was quantified using ezTrack (Pennington et al., 2021, 2019) in the first 10 seconds of the cue (cue window). To confirm that our findings are robust across thresholding decisions, we explored the following parameter spaces: minimum duration = 0.25s, 0.5s, 1s; motion threshold = 10px, 25px, 50px, 100px.

#### DeepLabCut

DeepLabCut (v2.2.1.1; (Lauer et al., 2022; Mathis et al., 2018; Nath et al., 2019) was used for markerless pose estimation. 95% of all labeled frames were used to train a ResNet-50 based neural network with default parameters for 500,000 training iterations. The train error was 4.48 pixels, and the test error was 5.84 pixels. The resulting neural network was used to track body parts for all cue-interval clips.

#### Keypoint-MoSeq

Keypoint-MoSeq (kp-MoSeq; v0.5.0; (Weinreb et al., 2024) was used for unsupervised behavioral classification. 16 cue clips for each cue type were semi-randomly selected from each session over 14 training and 1 recall session of the training mixed valence dataset. The model was fit over 50 iterations in the autoregressive only model and over an additional 450 iterations in the full model. Kappa was tuned to 1e5 for the autoregressive-only model and 5e4 for the full model to achieve a median syllable duration of 400ms as recommended by Weinreb et al. (2024).

kp-MoSeq uses a stochastic fitting procedure that will produce slightly different syllable segmentations when run multiple times with different random seeds. For this reason, 20 kp-MoSeq models were generated using 20 different seeds and the best model was selected based on maximal expected marginal likelihood.

#### Syllable post-processing

24 syllables were generated and manually labeled by an experimenter (*Table S1*) into one of the following categories to comprehensively profile behavior and support interpretability: Attend, jump, rear, locomote, turn, other (groom, pause, lick). Syllable cumulative duration (referred to as ‘duration’) was quantified as the sum of frames that a syllable was present during a given time window.

#### Location-based predictors

We included location-based predictors to incorporate spatially salient points in the prediction model. We reasoned that the food port and corners of the conditioning chambers define salient locations during reward seeking and threat anticipation, respectively. Distance to the food port was calculated as distance from snout to food port. Food port orientation was calculated as follows: distance from tail base to food port / distance from snout to food port. Thus, values above 1 indicate closer snout distance (i.e., port orientation), whereas values below 1 indicate closer tail base distance (i.e., orientation away from port). Distance of tail base to each corner was calculated and the smallest distance included as a predictor. All distances were measured in pixels.

### Statistical analysis

#### Linear mixed-effects regression (LMER)

Repeated measures data were modeled using linear mixed-effects regression. Mouse ID was added as a random effect in all models to account for interdependence of observations within individuals across days. Planned contrasts were performed to determine statistical significance of comparisons using each model’s estimated marginal means. Multiple comparisons were performed using Tukey’s HSD to control the family-wise error rate.

#### Partial least squares discriminant analysis (PLS-DA)

PLS-DA is a supervised statistical approach used for classification and dimensionality reduction (Rosipal and Krämer, 2006). This method performs well with large numbers of correlated predictors, as it inherently provides classification and feature selection. Here, we used PLS-DA to predict CS identity from syllable duration and location-based predictors on the recall day data. We used data from the discovery cohort as training data and data from a separate validation cohort as test data to assess prediction accuracy.

Model performance was evaluated for two-way and three-way classification problems (Fig. S1). Two-way classification consistently outperformed three-way classification, and we therefore used two-way classification models for all downstream analyses. We further compared performance of prediction using different cue windows and found that 5 seconds mid-cue performed best. In addition, we compared sex-specific classifiers to each other as well as to classifiers using data of both sexes combined (Fig. S1).

#### Software

All analyses were performed in R (v4.3.1). LMERs were fit using the *lme4* package (v1.1.35.5). Planned contrasts were performed using the *emmeans* package (v1.8.9). PLS-DA was performed using the *mdatools* package (v0.14.2). All data is available at https://osf.io/4xfz2/. All analysis code is available at https://github.com/heike-s/ValenceProfile.

## Results

### Standard metrics of valence learning suggest limited acquisition

Because valenced stimuli are inherently salient, single valence paradigms conflate salience and valence. To better isolate valence encoding, we developed a paradigm in which mice learn appetitive and aversive associations in parallel. In this mixed valence paradigm, one auditory stimulus was paired with chocolate milk (CS^R^), another auditory stimulus was paired with mild footshock (CS^S^) and a third auditory stimulus was non-reinforced (CS^−^) (Fig. 1A). We compared behaviour of male and female mice trained in the mixed paradigm to mice trained in either an appetitive-only paradigm (only CS^R^ and CS), or an aversive-only paradigm (only CS^S^ and CS^−^).

**Figure 1.**
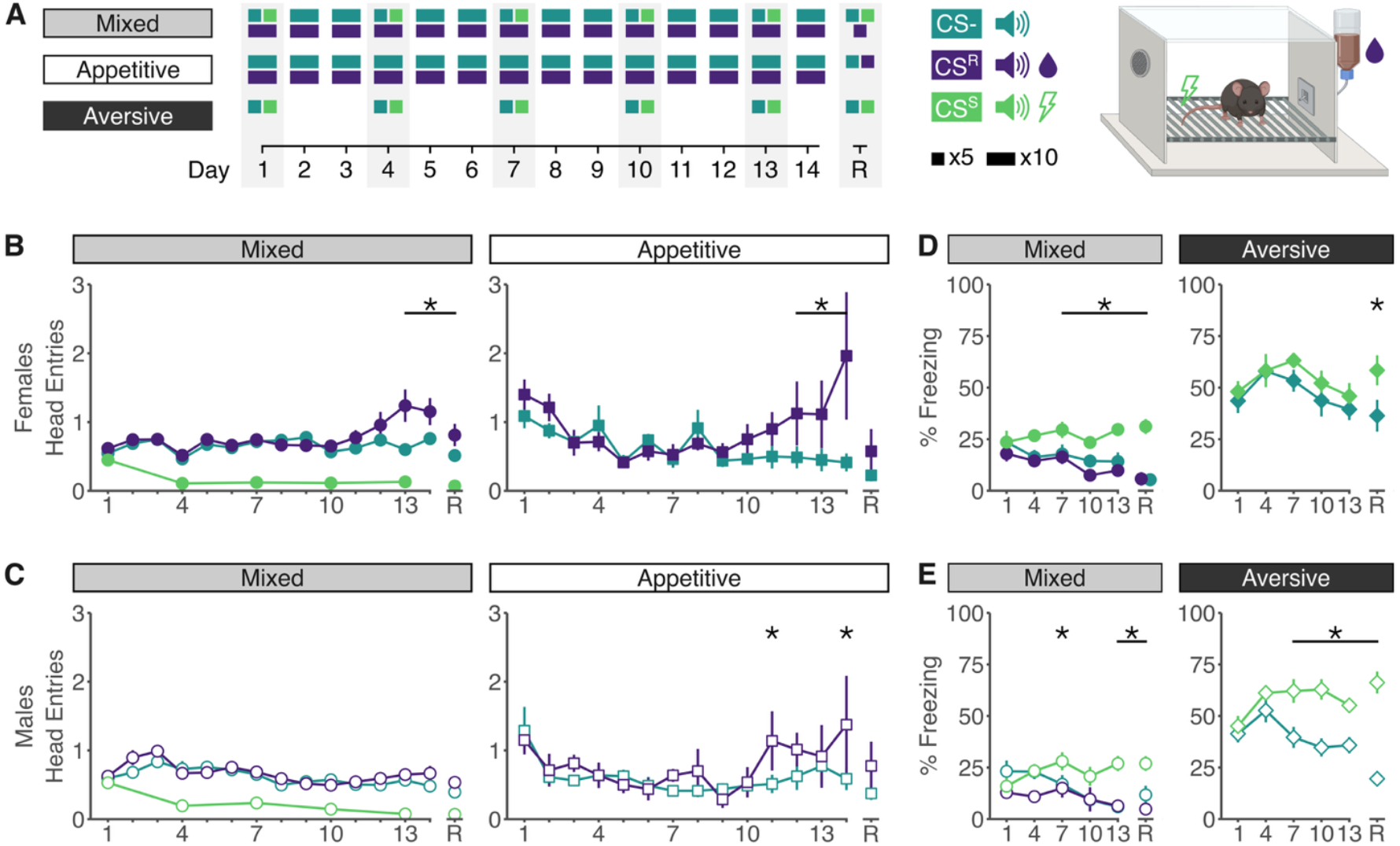
Standard indicators of valence learning point to limited acquisition. (**A**) Overview of the experimental design and apparatus. (**B**) In females, head entries in the first 10 seconds of the cue are significantly higher to the CS^R^ compared to the CS^−^ at the end of training in both mixed (left) and appetitive-only (right) paradigms. On recall day (R), discrimination in head entries only remains significant in the mixed paradigm. (**C**) Male mice are performing more head entries to the CS^R^ compared to the CS^−^ only at the end of the appetitive-only paradigm, which returns to non-significance on recall day. *Table S2* contains full statistical analyses, including comparison to CS^S^. (**D**) In females, percent time spent freezing is increased in response to CS^S^ compared to CS^−^ in the mixed valence paradigm, but not in the aversive only paradigm throughout training. On recall day, females have significantly higher freezing levels to the CS^S^ than the CS^−^ in both paradigms. (**E**) Male mice have increased CS^S^ freezing in the mixed and aversive-only paradigm as compared to CS^−^, and discrimination remains significant in both paradigms on recall day. *Table S3* contains full statistical analyses, including comparison to CS^R^. * *padj* < 0.05.

We first evaluated learning in all three conditioning paradigms using field standard metrics: number of head entries to evaluate appetitive learning and percent time freezing to evaluate aversive learning. Quantifying behavioral responding during the first 10 seconds of each cue (prior to outcome delivery) revealed limited evidence of appetitive acquisition in the mixed or appetitive-only paradigm, with significant increases in head entries to the CS^R^ only at training end in female (Fig. 1B) or male (Fig. 1C) mice. This is consistent across other food port metrics (Fig. S2). In contrast, we observed stronger evidence of CS^S^ acquisition, yet behavioural responding varied between males and females across paradigms (Fig. 1D,E). In the mixed paradigm, both males and females freeze more to the CS^S^ than either CS^−^ or CS^R^, whereas in the aversive-only paradigm males again freeze more to the CS^S^ than CS^−^ but females show high levels of freezing to both cues. These observations are robust across a range of freezing thresholds (Fig. S3). Further, overall freezing levels were higher in the aversive-only paradigm. Due to the lack of evidence for reward learning, as well as the paradigm differences in freezing, we asked if these standard behavioural metrics accurately capture behavioural expression of learning.

### Data-driven analysis identifies robust behavioural predictors of cue identity

Standard behavioural metrics quantify a predetermined subset of all behavioral variability. In contrast, data-driven methods enable comprehensive profiling of the full behavioral repertoire. Given increasing recognition that task parameters and sex can influence behavioural expression, we applied DeepLabCut (DLC; (Lauer et al., 2022; Mathis et al., 2018; Nath et al., 2019) and Keypoint-MoSeq (Weinreb et al., 2024) to agnostically identify and group recurrent behavioural patterns. We reasoned that if mice had indeed acquired cue-outcome associations, some aspect of behavioural patterns would predict cue type (CS^S^, CS^R^, CS^−^). To test this, we used PLS-DA regression to predict cue type from recall test behaviour (no outcomes delivered) (Fig. 2A). Two-way classifiers differentiating cue pairs in the mixed paradigm (CS^−^ *vs* CS^R^, CS^−^ *vs* CS^S^, CS^R^ *vs* CS^S^) achieved high classification accuracy in both male and female mice indicating that mice had indeed learned both appetitive and aversive associations (Fig. 2B,C). Applying the classifiers trained on the mixed paradigm to the single valence paradigms also yielded above chance classification accuracy, suggesting that mice express valence responses similarly across mixed and single valence paradigms.

**Figure 2.**
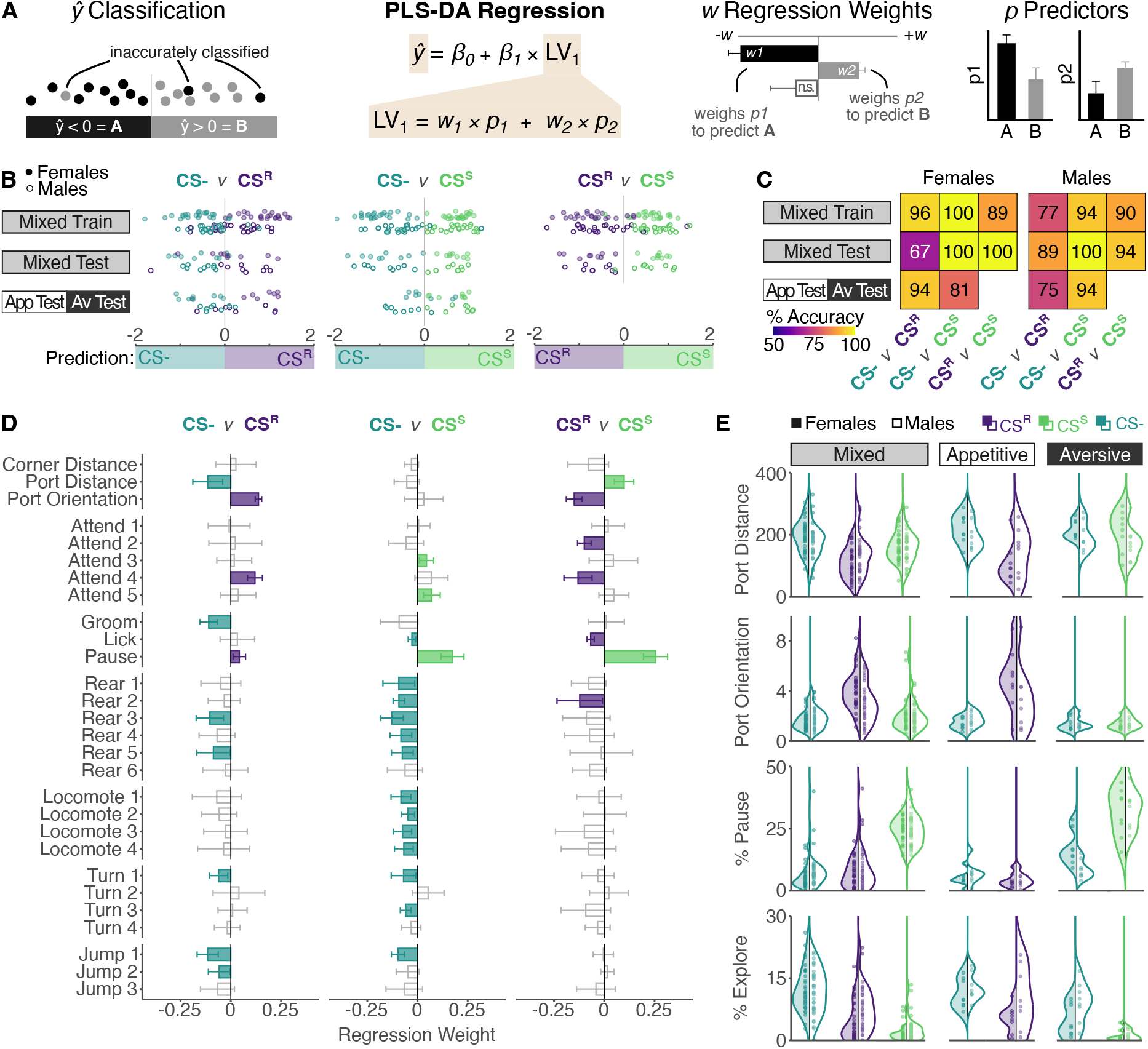
Cue identity can be reliably predicted from data-driven analysis approaches. (**A**) Overview of PLS-DA approach.(**B**) Predictions of cue-type on recall day using two-way classifiers in individual female (filled points) and male (empty points) mice in the mixed training dataset, mixed test dataset and appetitive- and aversive-only datasets. Point colors indicate true cue type, with predicted cue type indicated by position on x-axis. (**C**) Percent accuracy of cue type prediction in females (left) and males (right). All accuracy values surpass chance prediction (50%). (**D**) Regression weights for individual predictors for each two-way classifier. Significant predictors (*p*<0.05) are filled with the color of the cue type an increased value is predictive of, non-significant predictors are grey. (**E**) Distributions of variables predicting cue types. Port distance is decreased and port orientation is increased during the CS^R^, pausing is increased during the CS^S^, and exploration (sum of all rearing and locomoting syllables) is highest during the CS^−^.

We then asked which behavioral patterns contribute most strongly to cue type prediction by examining the regression weights of the PLS-DA regression latent variable. Food-port related location-based metrics, ‘*port-distance*’ and ‘*port-orientation*’, strongly contributed to predicting CS^R^ from CS^−^ (Fig. 2C). Increased *‘grooming’, ‘rearing’* and *‘locomoting’* behaviors contributed to predicting CS^−^ from CS^R^. ‘*Pausing*’ behaviour most strongly contributed to predicting CS^S^ from CS^−^ (Fig. 2C), and again CS^−^ associated with similar patterns of general movement. Differentiating CS^R^ from CS^S^ again relied on food-port related location-based metrics for CS^R^ and ‘*pausing’* for CS^S^ suggesting that these are useful behavior metrics to identify appetitive and aversive responses. Visualizing raw data for these behavioural predictors confirmed expected distributions across cue-types (Fig. 2D, Fig. S4). Overall, this suggests that the lack of evidence of acquisition observed with standard behavioural metrics reflects the limitations of the metrics in a mixed-valence setting rather than a lack of learning.

### Apparent sex differences in learning reflect underlying context-driven exploration

Having built robust behavioral classifiers, we applied these to probe how learning develops in male and female mice over training across paradigms (Fig. 3A). In the mixed paradigm, both males and females rapidly discriminate CS^R^ from CS^S^ indicating they quickly acquire valence-specific associations. In contrast, discriminating either valenced cue from the CS^−^ varied markedly across paradigms and sexes. By mid-training, females discriminate CS^R^ from CS^−^ in both the mixed and appetitive-only paradigm. In contrast, although males learn to discriminate the CS^R^ from CS^−^ in both paradigms, this occurs later than in females, appearing robustly only towards training end. Uniquely, in males in the appetitive-only paradigm, the CS^R^ is initially misclassified as CS^−^ suggesting an altered trajectory of CS^R^ versus CS^−^ discrimination. Notably, despite these differences, males eventually acquire the CS^R^ association and CS^−^ discrimination in both protocols (Fig. S5, Fig. 3B).

**Figure 3.**
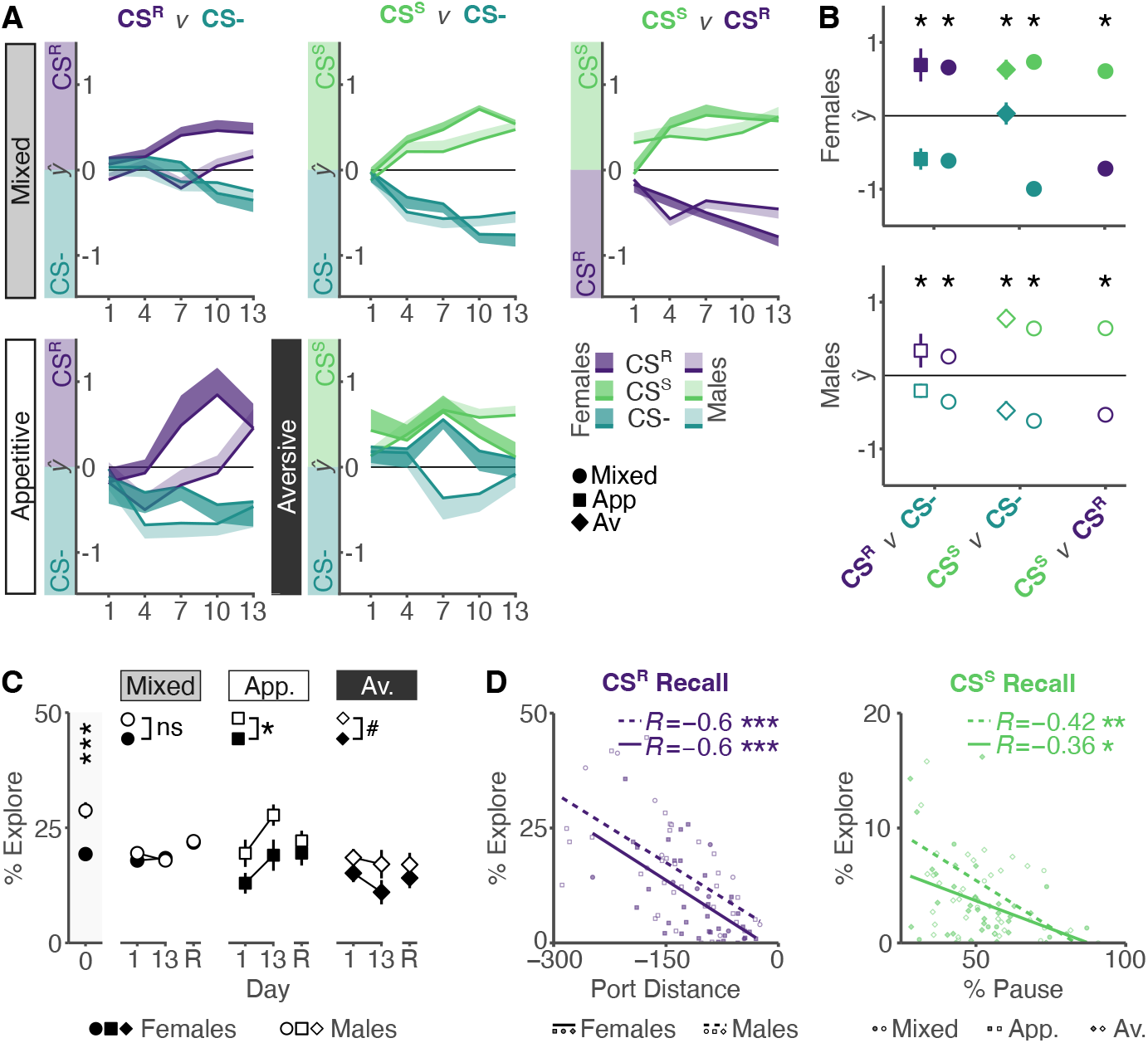
Learning trajectories are mediated by underlying sex differences in exploration. (**A**) Cue type prediction throughout training in the mixed paradigm (top) and single-valence paradigms (bottom) throughout training in each two-way classifier. Male and female predictions largely follow each other in the mixed paradigm, whereas sex-specific learning trajectories can be observed in both the appetitive and aversive-only paradigm. (**B**) Cue type prediction on recall day for each paradigm in females (top) and males (bottom) shows cue discrimination between all cue types in both sexes (<u>Contrast</u> CS^−^ – CS^R^: Appetitive female: *t-ratio_360_* = −4.635, *p_adj_*< 0.001; Mixed female: *t-ratio_360_*= −7.92, *p_adj_*< 0.001; Appetitive male: *t-ratio_360_*= −0.547, *p_adj_*< 0.05; Mixed male: *t-ratio_360_*= −3.629, *p_adj_*< 0.001; <u>Contrast CS^−^ – CS^S^</u>: Aversive female: *t-ratio_360_*= −2.165, *p_adj_*< 0.05; Mixed female: *t-ratio_360_*= −15.1, *p_adj_*< 0.001; Aversive male: *t-ratio_360_*= −4.544, *p_adj_*< 0.001; Mixed male: *t-ratio_360_*= −11.746, *p_adj_*< 0.001; <u>Contrast CS^R^ – CS^S^</u>: Mixed female: *t-ratio_360_*= −7.18, *p_adj_*< 0.001; Mixed male: *t-ratio_360_*= −8.117, *p_adj_*< 0.001). The full statistical model is reported in *Table S4.* (**C**) Percent time spent exploring in the pre-cue period during habituation, first day of training (day 1), end of training (day 13) and on recall test (R). Males and females differ in their levels of exploration on habituation (<u>Contrast female – male:</u> *t-ratio_30_*= −4.281, *p* < 0.001), and throughout training in the single-valence paradigms (<u>Contrast female – male:</u> Appetitive: *t-ratio_85.9_*= −2.559, *p* < 0.05; Aversive: *t-ratio_88.1_*= −1.784, *p* = 0.078) but not the mixed-valence paradigm (<u>Contrast female – male:</u> *t-ratio_94.3_*= −0.373, *p* = 0.71). The full statistical model is reported in *Table S5.* (**D**) Cue-relevant behaviors port distance (CS^R^ predictive, left) and pausing (CS^S^ predictive, right) negatively correlate with % exploration during cue presentation. * *p_adj_*< 0.05, ** *p_adj_*< 0.01, *** *p_adj_*< 0.001.

Intriguingly, predicting CS^S^ from CS^−^ revealed distinct effects of sex. Females rapidly discriminate CS^S^ from CS^−^ in the mixed paradigm but, in the aversive-only paradigm, the CS^−^ is consistently misclassified as CS^S^ throughout training, with discrimination only evident at recall (Fig. 3A,B). In contrast, males similarly discriminate CS^S^ from CS^−^ in both paradigms. The misclassification of CS^R^ as CS^−^ in males and CS^−^ as CS^S^ in females suggests that males are exhibiting more CS^−^ associated behaviours (‘*rearing’*, ‘*locomoting’*) during the CS^R^ and that females are showing less of these same behaviours during the CS^−^ and more of the CS^S^ associated *pausing* behaviours. Integrating these observations led us to hypothesize that a common underlying factor of exploration might explain the observed sex differences in cue responding in single valence protocols.

We reasoned that, if exploration drives sex-specific cue responding, sex differences in exploration should be apparent even before learning. Specifically, we predicted that males explore more than females. To test this, we compared exploration on the habituation day and pre-cue periods during training by quantifying the total percent time in any *rearing* or *locomoting* syllable (from here on summarized as ‘*exploring’;* Fig. 3C). This confirmed that males explore more than females throughout the habituation day when no valenced outcomes have been presented. During the pre-cue period there were no sex differences in exploration in the mixed valence paradigm. However, as predicted, in the appetitive-only paradigm, males explored more than females and in the aversive-only paradigm this trend was maintained. To confirm that elevated exploration inhibits the expression of valence-specific behaviours, we correlated these metrics, confirming the predicted negative relationships (Fig. 3D). Overall, this demonstrates that behavioural expression in single valence paradigms is biased by sex differences in exploration. This bias is not present in a mixed valenced paradigm, suggesting that a mixed valenced paradigm provides a more accurate assessment of valence learning. Performance in the mixed valenced paradigm demonstrates that both sexes *can* similarly learn valenced associations, yet it is not clear why females fail to do this in the aversive-only context. To understand why sex differences in exploration should be specific to this single valence paradigm we interrogated the specific conditions under which these sex differences are expressed.

### Shock exposure broadly facilitates *pausing* to suppress exploration in female mice

To gain insight into paradigm specific sex differences in exploration we compared exploration across the mixed-valence and aversive-only paradigms. We calculated the percent time spent in exploration or the relative absence of exploration, defined as ‘*pausing’*, in the pre-cue period to determine if context modulates behaviour independently of cues. In females, but not in males, levels of exploration and pausing differed between paradigms with overall higher exploration and lower pausing in the mixed paradigm (Fig. 4A). This suggests that the outcomes experienced during acquisition and the associative context modulate baseline exploration behaviour in females more strongly than in males.

**Figure 4.**
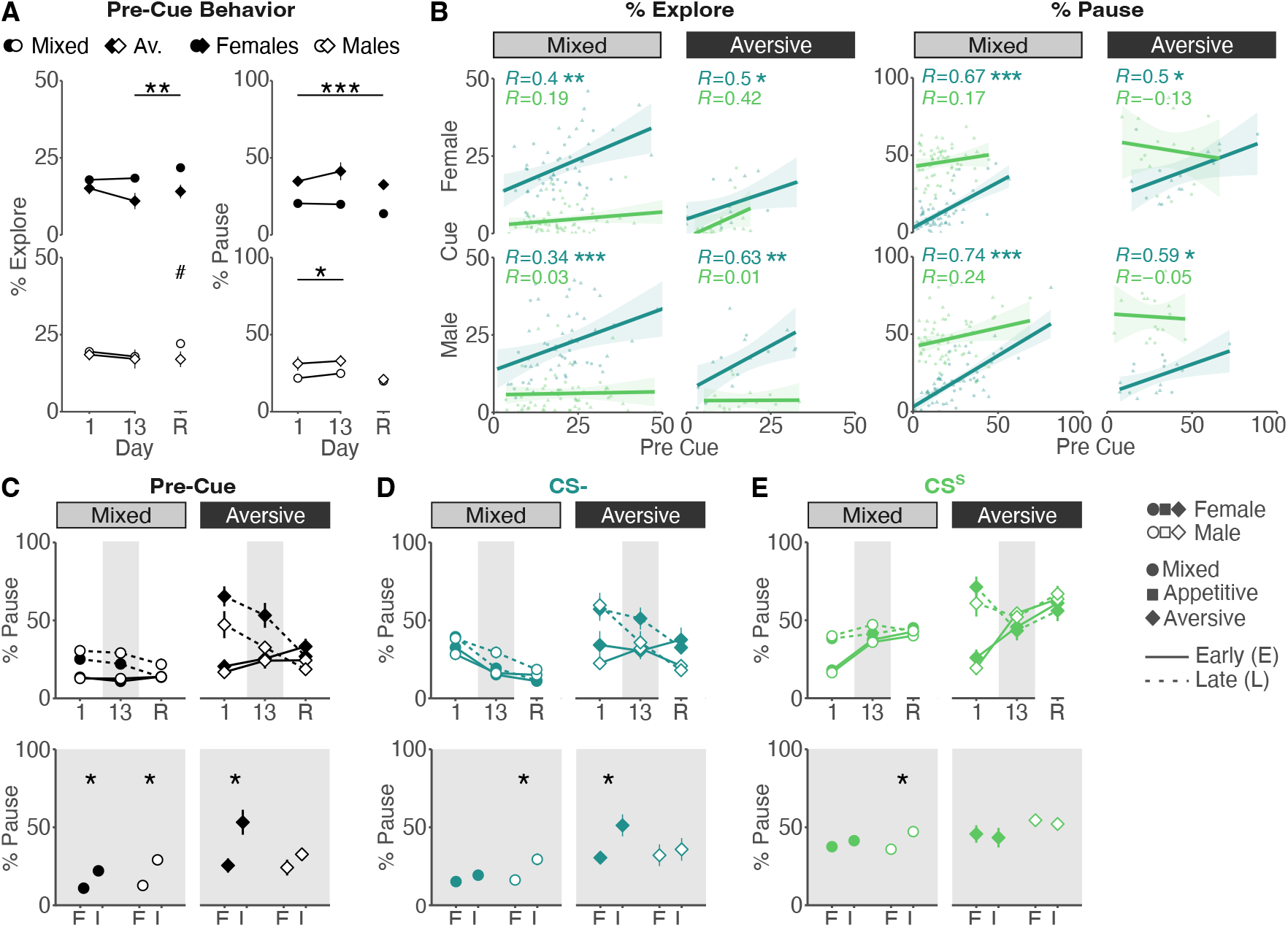
Repeated shock exposure promotes widespread pausing, limiting exploratory behavior in female mice. (**A**) Pre-cue exploration and pausing behaviors show significant differences between mixed valence and aversive-only paradigms throughout training and recall in females (top; <u>Contrasts explore Mixed *vs* Aversive</u>: all days: *t-ratio*_*77*.*5*_ = −6.201, *p*_*adj*_ < 0.001; day 1: *t-ratio*_*195*_ = −3.548, *p*_*adj*_ < 0.001; day 13: *t-ratio*_*191*_ = −5.219, *p*_*adj*_ < 0.001; day R: *t-ratio*_*186*_ = −4.81, *p*_*adj*_ < 0.001; <u>Contrasts pause Mixed *vs* Aversive</u>: all days: *t-ratio*_*77*_ = 3.338, *p*_*adj*_ < 0.01; day 1: *t-ratio*_*213*_ = 1.025, *p*_*adj*_ = 0.306; day 13: *t-ratio*_*212*_ = 2.7, *p*_*adj*_ < 0.01; day R: *t-ratio*_*210*_ = 2.886, *p*_*adj*_ < 0.01). In males (bottom), time spent exploring is comparable across paradigms, and time spent pausing is only modestly increased in the aversive-only paradigm during training (<u>Contrasts exploring Mixed *vs* Aversive</u>: all days: *t-ratio*_*73*.*6*_ = −2.159, *p*_*adj*_ < 0.05; day 1: *t-ratio*_*185*_ = −2.419, *p*_*adj*_ < 0.05; day 13: *t-ratio*_*187*_ = −2.051, *p*_*adj*_ < 0.05; day R: *t-ratio*_*185*_ = −0.288, *p*_*adj*_ = 0.774; <u>Contrasts pausing Mixed *vs* Aversive</u>: all days: *t-ratio*_*72*_ = 1.288, *p*_*adj*_ = 0.202; day 1: *t-ratio*_*210*_= 0.367, *p*_*adj*_ = 0.714; day 13: *t-ratio*_*210*_ = 0.277, *p*_*adj*_ = 0.782; day R: *t-ratio*_*210*_ = 1.911, *p*_*adj*_ = 0.057). The full statistical model is reported in *Table S4*. (**B**) Correlations between CS^−^ and CS^S^ pre-cue and cue exploration (left) and pausing (right) behaviors show that CS^−^ behaviors correlate significantly with pre-cue behaviors, whereas CSS behaviors do not. (**C**) Percent time spent pausing increases over the course of the session from the first two cues (early; E) to the last two cues (late; L) in the pre-cue period on the first day of training in both paradigms and sexes (<u>Contrasts day 1 early *vs* late</u>: females mixed: *t-ratio*_*360*.*7*_ = −3.381, *p*_*adj*_ < 0.001; females aversive: *t-ratio*_*360*.*7*_ = −6.097, *p*_*adj*_ < 0.001; males mixed: *t-ratio*_*360*.*7*_ = −5.249, *p*_*adj*_ < 0.001; males aversive: *t-ratio*_*360*.*7*_ = −4.424, *p*_*adj*_ < 0.001). On day 13 (resolved in bottom plot), both male and female mice show small but significant increases in pausing at the end of the session while, in the aversive-only paradigm, females show a large increase in pausing at the end of the session absent in males (<u>Contrasts day 13 early *vs* late</u>: females mixed: *t-ratio*_*360*.*7*_ = −2.802, *p*_*adj*_ < 0.01; females aversive: *t-ratio*_*360*.*7*_ = −4.023, *p*_*adj*_ < 0.001; males mixed: *t-ratio*_*360*.*7*_ = −4.613, *p*_*adj*_ < 0.001; males aversive: *t-ratio*_*360*.*7*_ = −1.229, *p*_*adj*_ *=* 0.22). On recall day (R), pausing levels are only significantly elevated at the end of the session in males in the mixed paradigm (<u>Contrasts day R early *vs* late</u>: females mixed: *t-ratio*_*360*.*7*_ = 0.113, *p*_*adj*_ = 0.91; females aversive: *t-ratio*_*360*.*7*_ = 0.921, *p*_*adj*_ = 0.353; males mixed: *t-ratio*_*360*.*7*_ = −2.355, *p*_*adj*_ < 0.05; males aversive: *t-ratio*_*360*.*7*_ = 0.832, *p*_*adj*_ *=* 0.406). (**D**) Similarly, percent time spent pausing to the CS^−^ increases throughout the session on day 13 in females in the aversive-only paradigm, with a more modest increase in males in the mixed-valence paradigm (<u>Contrasts day 13 early *vs* late</u>: females mixed: *t-ratio*_*356*.*5*_ = −0.925, *p*_*adj*_ = 0.355; females aversive: *t-ratio*_*356*.*5*_ = −2.694, *p*_*adj*_ < 0.01; males mixed: *t-ratio*_*360*.*7*_ = −3.332, *p*_*adj*_ < 0.001; males aversive: *t-ratio*_*360*.*7*_ = −0.492, *p*_*adj*_ *=* 0.623). (**E**) On day 13 levels of pausing are stable from early to late session, except for a modest increase in males in the mixed paradigm (<u>Contrasts day 13 early *vs* late</u>: females mixed: *t-ratio*_*356*.*7*_ = −0.96, *p*_*adj*_ = 0.338; females aversive: *t-ratio*_*356*.*7*_ = 0.335, *p*_*adj*_ = 0.738; males mixed: *t-ratio*_*357*.*9*_ = −3.164, *p*_*adj*_ < 0.01; males aversive: *t-ratio*_*356*.*7*_ = 0.347, *p*_*adj*_ *=* 0.729). Significance levels are shown for day 13 comparisons only (bottom plots), with full statistical results reported in *Table S5*. ^#^ *p*_*adj*_ < 0.1, * *p*_*adj*_ < 0.05, ** *p*_*adj*_ < 0.01, *** *p*_*adj*_ < 0.001.

In the aversive-only paradigm, we demonstrated that high levels of ‘*pausing’* to CS^−^ obscure CS^S^ discrimination (Fig. 3A). Having now shown that high levels of pausing are also observed in the pre-cue period, we asked if the elevated freezing to CS^−^ is cue-driven or reflects a continuation of elevated context-mediated freezing. To test this, we correlated *exploration* and *pausing* between the pre-cue and cue periods for CS^−^ and CS^S^. We reasoned that, if cue onset elicits a distinct behavioural response, behaviour between pre-cue and cue periods will not correlate. Indeed, for the CS^S^, we observed no significant pre-cue to cue correlation in either paradigm or sex (Fig. 4B). However, for the CS^−^, we observed consistent, strong correlation across paradigms and sexes indicating that behaviour during the CS^−^ cue period is a continuation of contextually mediated behaviour and not a distinct cue-mediated response.

Altered pre-cue responding likely reflects the combination of the acute impact of innately appetitive or aversive outcomes encountered during the current session and the evolving associative history of the context. Both factors contribute on training days, whereas on the recall day, only the associative history of the context is present. In the aversive only paradigm, while females failed to show evidence of CS^−^ vs CS^S^ discrimination during training, this was clearly evidenced on the recall day (Fig. 3B). We therefore hypothesized that pre-cue differences are in part attributable to acute effects of outcome presentations. We reasoned that this effect will be amplified across the session as the impact of successive outcomes accrues but would not be apparent on the non-reinforced recall session. To test this, we compared *pausing* during the pre-cue, CS^S^ and CS^−^ periods for early (first two) and late (final two) cue presentations on training days 1 and 13 and recall (Fig. 4C-E). In the mixed paradigm, as expected, male and female mice *paused* more in the late pre-cue periods on both training day 1 and 13 but not during recall, suggesting a modest influence of outcomes. In the aversive-only paradigm, this modulation was stronger in both sexes on training day 1 however, by training end, this was no longer evident in males, but females continued to show elevated *pausing* across the session (Fig. 4C). This suggests that males, but not females, habituated to footshock, downregulating outcome-induced pausing across training days. As predicted, in females, this phenomenon was also evident in CS^−^ responding (Fig. 4D) but no such difference was observed for the CS^S^ (Fig. 4E). Overall, this suggests that, in females shock exposure acutely potentiates *pausing* specifically in the aversive only paradigm and that this obscures CS^−^ discrimination. This demonstrates that task-dependent sex differences in non-associative learning processes can mask similar underlying associative learning.

## Discussion

Despite the fundamental importance of valence learning to health and disease, limited research has simultaneously explored associative valence learning in male and female mice. Here, we compared mixed-valence and single-valence paradigms and revealed critical effects of task design on exploration and habituation, a form of non-associative learning, that interact with sex to alter behavioral expressions of learning. We demonstrate that both male and female mice can successfully acquire appetitive and aversive cue-outcome associations across mixed and single-valence tasks. Using data-driven approaches to interrogate cue-mediated behavioral changes, we identify robust metrics of acquisition of cue-outcome associations that are applicable across paradigms and sexes. Critically, we show that behavior in single-valence paradigms is biased by pre-existing sex differences in exploratory behaviors that mask behavioral expression of cue learning, potentially leading to erroneous conclusions of sex differences in associative valence learning. Our analyses highlight the superiority of mixed-valence paradigms in minimizing baseline sex differences in exploratory behaviors, thereby allowing more accurate assessment of learning.

Our findings shed new light on previously reported sex differences in single valence learning tasks. Fear conditioning studies report stronger threat generalization and higher levels of conditioned responding in female rodents (Day et al., 2020, Day et al., 2016; Olivera-Pasilio and Dabrowska, 2023; Trott et al., 2022). Superficially, we replicate these findings in an aversive-only protocol, but comparison to the mixed-valence protocol suggests that these observations do not reflect an inherent inability of female mice to discriminate between the CS^S^ and the CS^−^. Rather, we show that the lack of discrimination is driven by lack of habituation to repeated shock, a form of non-associative learning (Ardiel and Rankin, 2010), and pre-existing sex differences in the propensity to engage in exploration. Research in rats has previously shown darting as a female-specific sexually divergent response in fear conditioning, a feature we were not able to identify in our datasets, possibly due to species or apparatus differences (Gruene et al., 2015; Mitchell et al., 2024, Mitchell et al., 2022). Research in reward learning, while sparser, has also suggested sex differences, with male rodents showing less robust reward conditioning (Lefner et al., 2022). Here, we demonstrate that apparent sex differences in single valence learning tasks are driven by underlying differences in exploration that mask cue learning in each sex depending on context and are not true learning differences.

Sex differences in exploratory behavior in mice have previously been reported in other tasks. For example, in a reward-based decision-making task, males had overall higher levels of exploration than females, but females were able to learn from exploration more quickly (Cathy S Chen et al., 2021). Curiously, other research suggests increased velocity or rearing in female rodents compared to males in the open field test or social interaction test (McElroy and Howland, 2025; Schuler et al., 2024). This indicates that sex differences in exploration may be task specific with increased exploration in females in anxiety-inducing contexts, consistent with reports that increased locomotion is a female-specific anxiety-like behavior (Gruene et al., 2015; Mitchell et al., 2022; Schuler et al., 2024). Overall, this study adds to the growing literature on the nuances of sex differences in behavior (Cathy S. Chen et al., 2021; Greiner et al., 2019; Gruene et al., 2015; Pellman et al., 2017; Shansky, 2018; Tronson, 2018).

Our analysis highlights the importance of task design in shaping behavioral expression. We find elevated periods of immobility in the aversive-only paradigm compared to the mixed-valence paradigm for both sexes, suggesting an increased influence of contextual learning in single-valence fear learning. This is consistent with the varying predictive value of contexts between single and mixed-valence paradigms. In the single-valence paradigm, the only salient outcome that is encountered is foot-shock, making the context a relatively good predictor of outcome. In contrast, in the mixed-valence paradigm, two salient, oppositely valenced outcomes are encountered – footshock and reward-rendering the context a poor predictor of outcome. In this way, mixed-valence contexts reduce contextual learning and better isolate cue-modulation of behavior, limiting the influence of pre-existing sex differences in exploration on behavioral expression. Reduced habituation to repeated shock in female mice was also task dependent and was restricted to the aversive-only setting. This suggests a female-specific sensitivity to repetitive aversive experiences, suggesting a potentially important sex-specific phenotype for future studies with relevance to human sex differences in neuropsychiatric conditions. Our findings emphasize the need for caution in interpreting sex differences in task performance and illustrate the impact of task design. In line with previous research, our findings also emphasize the power of rigorously applied data-driven approaches in behavioral neuroscience in re-evaluating long-standing assumptions and identifying sex-specific phenotypes in behaviors previously established on male data only (Levy et al., 2023; Schuler et al., 2024; Shansky, 2024).

We established a mixed-valence paradigm with robust behavioral readouts in male and female mice. While similar tasks have been previously designed in rats or in mouse head-fixed setups, it has been challenging to study approach and avoidance behavior in a within-subject design in freely moving mice. We note that, specifically for approach behavior, standard readouts such as head entries or licks perform poorly in quantifying individual learning. This challenge is perhaps reflected in the scarcity of publications that report cue-based reward learning in freely moving mice. We observed that in both appetitive-only and mixed-valence paradigms, mice do not physically enter the food port until reward delivery yet other approach behaviors are evident in video-based quantification. Using advanced behavioral classification to overcome this challenge, we identified simple behavioral measures that can be easily applied in future using simpler tracking methods (e.g., ezTrack). The mixed-valence paradigm effectively isolates valence learning and overcomes limitations of single-valence approaches to model real-world learning in which mixed valence outcomes are commonly encountered. However, several important task parameters that influence learning and behavior remain to be further explored, including varying the US modality and US salience (Mitchell et al., 2024). Further, the precise setup of the conditioning apparatus will likely also influence task performance. For example, our use of a non-retractable food port allowed mice to consume chocolate milk even after the cue ends, potentially slowing learning compared to a retractable reward delivery apparatus, in which mice can only consume chocolate milk during the cue period. We present a blueprint for rigorously identifying relevant behavioral metrics of learning in both sexes that can be applied across a wide variety of paradigms.

To conclude, we demonstrate that both males and females are able to acquire appetitive and aversive associations in both single and mixed-valence paradigms, yet the expression of this learning is shaped by task design. Our findings emphasize the critical importance of robust behavioral phenotyping in both sexes to achieve accurate insight into learning. We compared single and mixed-valence paradigms and revealed how task variables and experimental design considerations differentially impact behavioral expression in male and female mice in cued valence learning. Most importantly, we show that learning assessed in single valence paradigms is biased by sex-differences in exploration and that mixed-valence contexts minimize these non-task relevant sex differences to provide an unobscured assessment of learning in both sexes. Leveraging mixed valence learning protocols in combination with in vivo circuit interrogation techniques and animal models for disease promises to provide clear insight into the mechanisms of valence processing in health and disease.

## References

Admon R, Pizzagalli DA. 2015. Dysfunctional reward processing in depression. Curr Opin Psychol 4:114–118. doi:10.1016/j.copsyc.2014.12.011

Adolphs R, Mlodinow L, Barrett LF. 2019. What is an emotion? Curr Biol 29:R1060–R1064. doi:10.1016/j.cub.2019.09.008

Anderson DJ, Adolphs R. 2014. A Framework for Studying Emotions across Species. Cell 157:187–200. doi:10.1016/j.cell.2014.03.003

Ardiel EL, Rankin CH. 2010. An elegant mind: Learning and memory in Caenorhabditis elegans. Learn Mem 17:191– 201. doi:10.1101/lm.960510

Borkar CD, Dorofeikova M, Le Q-SE, Vutukuri R, Vo C, Hereford D, Resendez A, Basavanhalli S, Sifnugel N, Fadok JP. 2020. Sex differences in behavioral responses during a conditioned flight paradigm. Behav Brain Res 389:112623. doi:10.1016/j.bbr.2020.112623

Brittlebank AD, Scott J, Mark J, Williams G, Ferrier IN. 1993. Autobiographical Memory in Depression: State or Trait Marker? Br J Psychiatry 162:118–121. doi:10.1192/bjp.162.1.118

Carroll JN, Myers B, Vaaga CE. 2024. Repeated presentation of visual threats drives innate fear habituation and is modulated by environmental and physiological factors. bioRxiv 2024.10.15.618513. doi:10.1101/2024.10.15.618513

Chen Cathy S., Ebitz RB, Bindas SR, Redish AD, Hayden BY, Grissom NM. 2021. Divergent Strategies for Learning in Males and Females. Curr Biol 31:39-50.e4. doi:10.1016/j.cub.2020.09.075

Chen Cathy S, Knep E, Han A, Ebitz RB, Grissom NM. 2021. Sex differences in learning from exploration. eLife 10:e69748. doi:10.7554/elife.69748

Chiara GD. 1999. Drug addiction as dopamine-dependent associative learning disorder. Eur J Pharmacol 375:13– 30. doi:10.1016/s0014-2999(99)00372-6

Day HLL, Reed MM, Stevenson CW. 2016. Sex differences in discriminating between cues predicting threat and safety. Neurobiol Learn Mem 133:196–203. doi:10.1016/j.nlm.2016.07.014

Day HLL, Suwansawang S, Halliday DM, Stevenson CW. 2020. Sex differences in auditory fear discrimination are associated with altered medial prefrontal cortex function. Sci Rep 10:6300. doi:10.1038/s41598-020-63405-w

Deseyve C, Domingues AV, Carvalho TTA, Armada G, Correia R, Vieitas-Gaspar N, Wezik M, Pinto L, Sousa N, Coimbra B, Rodrigues AJ, Soares-Cunha C. 2024. Nucleus accumbens neurons dynamically respond to appetitive and aversive associative learning. J Neurochem 168:312–327. doi:10.1111/jnc.16063

Domingues AV, Carvalho TTA, Martins GJ, Correia R, Coimbra B, Bastos-Gonçalves R, Wezik M, Gaspar R, Pinto L, Sousa N, Costa RM, Soares-Cunha C, Rodrigues AJ. 2025. Dynamic representation of appetitive and aversive stimuli in nucleus accumbens shell D1-and D2-medium spiny neurons. Nat Commun 16:59. doi:10.1038/s41467-024-55269-9

Douglas KM, Porter RJ. 2010. Recognition of disgusted facial expressions in severe depression. Br J Psychiatry 197:156–157. doi:10.1192/bjp.bp.110.078113

Ehlers A, Clark DM. 2000. A cognitive model of posttraumatic stress disorder. Behav Res Ther 38:319–345. doi:10.1016/s0005-7967(99)00123-0

Elliott R, Sahakian BJ, Herrod JJ, Robbins TW, Paykel ES. 1997. Abnormal response to negative feedback in unipolar depression: evidence for a diagnosis specific impairment. J Neurol, Neurosurg Psychiatry 63:74–82. doi:10.1136/jnnp.63.1.74

Greiner EM, Müller I, Norris MR, Ng KH, Sangha S. 2019. Sex differences in fear regulation and reward-seeking behaviors in a fear-safety-reward discrimination task. Behav Brain Res 368:111903. doi:10.1016/j.bbr.2019.111903

Gruene TM, Flick K, Stefano A, Shea SD, Shansky RM. 2015. Sexually divergent expression of active and passive conditioned fear responses in rats. eLife 4:e11352. doi:10.7554/elife.11352

Gündem D, Potočnik J, Winter F-LD, Kaddouri AE, Stam D, Peeters R, Emsell L, Sunaert S, Oudenhove LV, Vandenbulcke M, Barrett LF, Stock JV den. 2022. The neurobiological basis of affect is consistent with psychological construction theory and shares a common neural basis across emotional categories. Commun Biol 5:1354. doi:10.1038/s42003-022-04324-6

Iordanova MD, Yau JO-Y, McDannald MA, Corbit LH. 2021. Neural substrates of appetitive and aversive prediction error. Neurosci Biobehav Rev 123:337–351. doi:10.1016/j.neubiorev.2020.10.029

Keiser AA, Turnbull LM, Darian MA, Feldman DE, Song I, Tronson NC. 2017. Sex Differences in Context Fear Generalization and Recruitment of Hippocampus and Amygdala during Retrieval. Neuropsychopharmacology 42:397–407. doi:10.1038/npp.2016.174

Konorski J. 1973. On two types of conditional reflex: General laws of association. Cond Reflex Pavlovian J Res Ther 8:2–9. doi:10.1007/bf03000279

Kuehner C. 2003. Gender differences in unipolar depression: an update of epidemiological findings and possible explanations. Acta Psychiatr Scand 108:163–174. doi:10.1034/j.1600-0447.2003.00204.x

Lauer J, Zhou M, Ye S, Menegas W, Schneider S, Nath T, Rahman MM, Santo VD, Soberanes D, Feng G, Murthy VN, Lauder G, Dulac C, Mathis MW, Mathis A. 2022. Multi-animal pose estimation, identification and tracking with DeepLabCut. Nat Methods 19:496–504. doi:10.1038/s41592-022-01443-0

Laurent V, Westbrook RF, Balleine BW. 2022. Affective Valence Regulates Associative Competition in Pavlovian Conditioning. Front Behav Neurosci 16:801474. doi:10.3389/fnbeh.2022.801474

Lee YI, Lee D, Kim H, Kim MJ, Jeong H, Kim D, Glotzbach-Schoon E, Choi S-H. 2024. Overgeneralization of conditioned fear in patients with social anxiety disorder. Front Psychiatry 15:1415135. doi:10.3389/fpsyt.2024.1415135

Lefner MJ, Dejeux MI, Wanat MJ. 2022. Sex Differences in Behavioral Responding and Dopamine Release during Pavlovian Learning. eNeuro 9:ENEURO.0050-22.2022. doi:10.1523/eneuro.0050-22.2022

Lefner MJ, Moghaddam B. 2024. Reward and punishment contingency shifting reveals distinct roles for VTA dopamine and GABA neurons in behavioral flexibility. bioRxiv 2024.10.07.617060. doi:10.1101/2024.10.07.617060

Levy DR, Hunter N, Lin S, Robinson EM, Gillis W, Conlin EB, Anyoha R, Shansky RM, Datta SR. 2023. Mouse spontaneous behavior reflects individual variation rather than estrous state. Curr Biol 33:1358-1364.e4. doi:10.1016/j.cub.2023.02.035

Lissek S, Kaczkurkin AN, Rabin S, Geraci M, Pine DS, Grillon C. 2014. Generalized Anxiety Disorder Is Associated With Overgeneralization of Classically Conditioned Fear. Biol Psychiatry 75:909–915. doi:10.1016/j.biopsych.2013.07.025

Lissek S, Powers AS, McClure EB, Phelps EA, Woldehawariat G, Grillon C, Pine DS. 2005. Classical fear conditioning in the anxiety disorders: a meta-analysis. Behav Res Ther 43:1391–1424. doi:10.1016/j.brat.2004.10.007

Maniglio R, Gusciglio F, Lofrese V, Murri MB, Tamburello A, Innamorati M. 2014. Biased processing of neutral facial expressions is associated with depressive symptoms and suicide ideation in individuals at risk for major depression due to affective temperaments. Compr Psychiatry 55:518–525. doi:10.1016/j.comppsych.2013.10.008

Mathis A, Mamidanna P, Cury KM, Abe T, Murthy VN, Mathis MW, Bethge M. 2018. DeepLabCut: markerless pose estimation of user-defined body parts with deep learning. Nat Neurosci 21:1281–1289. doi:10.1038/s41593-018-0209-y

McElroy DL, Howland JG. 2025. Sex differences in exploratory behavior of rats successfully performing the object-in-place recognition memory test. Behav Brain Res 477:115303. doi:10.1016/j.bbr.2024.115303

McLean CP, Asnaani A, Litz BT, Hofmann SG. 2011. Gender differences in anxiety disorders: Prevalence, course of illness, comorbidity and burden of illness. J Psychiatr Res 45:1027–1035. doi:10.1016/j.jpsychires.2011.03.006

Mitchell JR, Trettel SG, Li AJ, Wasielewski S, Huckleberry KA, Fanikos M, Golden E, Laine MA, Shansky RM. 2022. Darting across space and time: parametric modulators of sex-biased conditioned fear responses. Learn Mem 29:171–180. doi:10.1101/lm.053587.122

Mitchell JR, Vincelette L, Tuberman S, Sheppard V, Bergeron E, Calitri R, Clark R, Cody C, Kannan A, Keith J, Parakoyi A, Pikus M, Vance V, Ziane L, Brenhouse H, Laine MA, Shansky RM. 2024. Behavioral and neural correlates of diverse conditioned fear responses in male and female rats. Neurobiol Stress 33:100675. doi:10.1016/j.ynstr.2024.100675

Nath T, Mathis A, Chen AC, Patel A, Bethge M, Mathis MW. 2019. Using DeepLabCut for 3D markerless pose estimation across species and behaviors. Nat Protoc 14:2152–2176. doi:10.1038/s41596-019-0176-0

Noworyta K, Cieslik A, Rygula R. 2021. Neuromolecular Underpinnings of Negative Cognitive Bias in Depression. Cells 10:3157. doi:10.3390/cells10113157

Olivera-Pasilio V, Dabrowska J. 2023. Fear-conditioning to unpredictable threats reveals sex differences in rat fear-potentiated startle (FPS). bioRxiv 2023.03.06.531430. doi:10.1101/2023.03.06.531430

Pellman BA, Schuessler BP, Tellakat M, Kim JJ. 2017. Sexually Dimorphic Risk Mitigation Strategies in Rats. eNeuro 4:ENEURO.0288-16.2017. doi:10.1523/eneuro.0288-16.2017

Pennington ZT, Diego KS, Francisco TR, LaBanca AR, Lamsifer SI, Liobimova O, Shuman T, Cai DJ. 2021. ezTrack— A Step-by-Step Guide to Behavior Tracking. Curr Protoc 1:e255. doi:10.1002/cpz1.255

Pennington ZT, Dong Z, Feng Y, Vetere LM, Page-Harley L, Shuman T, Cai DJ. 2019. ezTrack: An open-source video analysis pipeline for the investigation of animal behavior. Sci Rep-uk 9:19979. doi:10.1038/s41598-019-56408-9

Ray MH, Moaddab M, McDannald MA. 2022. Threat and Bidirectional Valence Signaling in the Nucleus Accumbens Core. J Neurosci 42:817–833. doi:10.1523/jneurosci.1107-21.2021

Ray MH, Russ AN, Walker RA, McDannald MA. 2020. The Nucleus Accumbens Core is Necessary to Scale Fear to Degree of Threat. J Neurosci 40:4750–4760. doi:10.1523/jneurosci.0299-20.2020

Rosipal R, Krämer N. 2006. Subspace, Latent Structure and Feature Selection, Statistical and Optimization Perspectives Workshop, SLSFS 2005, Bohinj, Slovenia, February 23-25, 2005, Revised Selected Papers 34–51. doi:10.1007/11752790_2

Schuler H, Eid RS, Wu S, Tse Y-C, Cvetkovska V, Lopez J, Quinn R, Zhou D, Meccia J, Dion-Albert L, Bennett SN, Newman EL, Trainor BC, Peă CJ, Menard C, Bagot RC. 2024. Data-driven analysis identifies novel modulation of social behavior in female mice witnessing chronic social defeat stress. Biol Psychiatry. doi:10.1016/j.biopsych.2024.11.017

Shansky RM. 2024. Behavioral neuroscience’s inevitable SABV growing pains. Trends Neurosci 47:669–676. doi:10.1016/j.tins.2024.06.007

Shansky RM. 2018. Sex differences in behavioral strategies: avoiding interpretational pitfalls. Curr Opin Neurobiol 49:95–98. doi:10.1016/j.conb.2018.01.007

Singh S, Bermudez-Contreras E, Nazari M, Sutherland RJ, Mohajerani MH. 2019. Low-cost solution for rodent home-cage behaviour monitoring. PLoS ONE 14:e0220751. doi:10.1371/journal.pone.0220751

Tafreshiha A, Burg SA van der, Smits K, Bl?mer LA, Heimel JA. 2021. Visual stimulus-specific habituation of innate defensive behaviour in mice. J Exp Biol 224:jeb230433. doi:10.1242/jeb.230433

Tronson NC. 2018. Focus on females: a less biased approach for studying strategies and mechanisms of memory. Curr Opin Behav Sci 23:92–97. doi:10.1016/j.cobeha.2018.04.005

Trott JM, Krasne FB, Fanselow MS. 2022. Sex differences in contextual fear learning and generalization: a behavioral and computational analysis of hippocampal functioning. Learn Mem 29:283–296. doi:10.1101/lm.053515.121

Vandael K, Zaman J, Vervliet B. 2025. The Effect of Avoidance Behavior on Generalization of Threat-Expectancy. Collabra: Psychol 11. doi:10.1525/collabra.129192

Weinreb C, Pearl JE, Lin S, Osman MAM, Zhang L, Annapragada S, Conlin E, Hoffmann R, Makowska S, Gillis WF, Jay M, Ye S, Mathis A, Mathis MW, Pereira T, Linderman SW, Datta SR. 2024. Keypoint-MoSeq: parsing behavior by linking point tracking to pose dynamics. Nat Methods 21:1329–1339. doi:10.1038/s41592-024-02318-2

