## Supplementary Figures for "Sex-specific exploration accounts for differences in valence learning in male and female mice"

*Figure S1*. Model fitting and latent variable selection.
*Figure S2*. Port-based readouts of reward learning.
*Figure S3*. Freezing quantification using different parameters.
*Figure S4*. Keypoint-MoSeq behavioral syllable duration raw values.
*Figure S5*. $CS^R$ predictive behaviors in females and males.

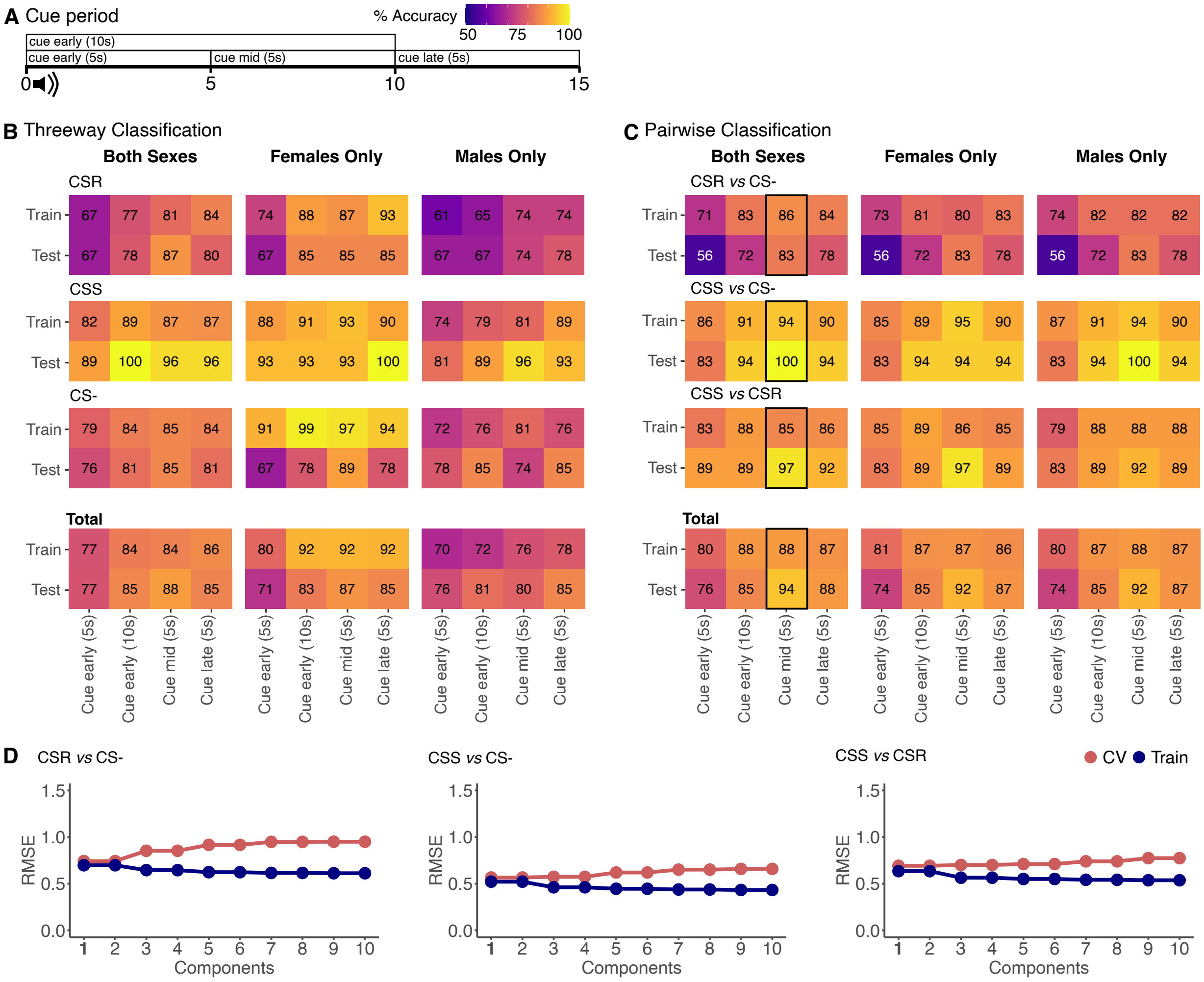


*Figure S1. Prediction accuracy and component (latent variable) selection.* (**A**) The cue period was split into 5 or 10 second intervals prior to predicting cue type identity. (**B**) Accuracy of cue type predictions in training and test dataset of three-way classifiers fitted using all data (left column), female only data (middle column) and male data (right column) in different using data from different cue intervals. (**C**) Same as B, but pairwise classifiers were trained to compare each cue type to the other. Best performance that generalizes to the test dataset was observed in the cue mid (5s) period using data from both sexes (values are enclosed in black boxes). (**D**) For each pairwise classifier, the optimal number of components (latent variables) is 1.

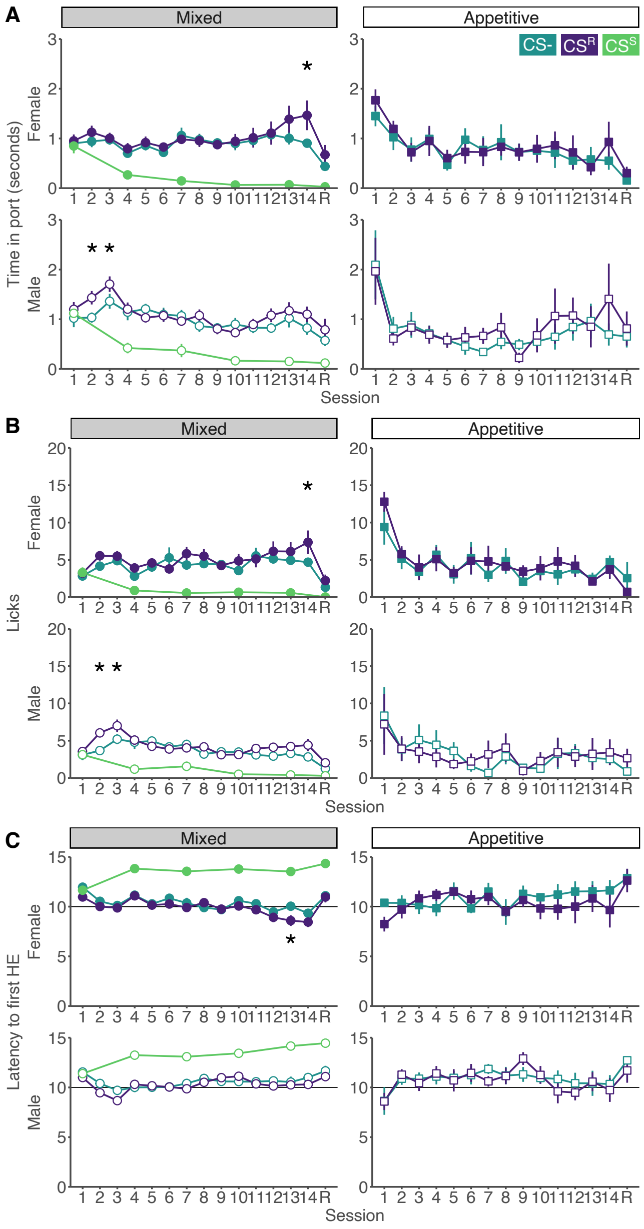


*Figure S2. Port-based readouts of reward learning.* (**A**) Time in port, (**B**) licks, and (**C**) latency to enter the food port during the cue in the mixed (left) and appetitive-only paradigms (right) in females (top) and males (bottom). Stars indicate significance between the $CS^R$ and $CS^-$, with full statistical results, including comparisons to the $CS^S$, reported in *Table S2*. Similar to head entries (Main Fig.1), other port-related metrics do not show evidence of robust associative reward learning in either the mixed or the appetitive-only paradigm in either sex. * $p_{adj} < 0.05$.

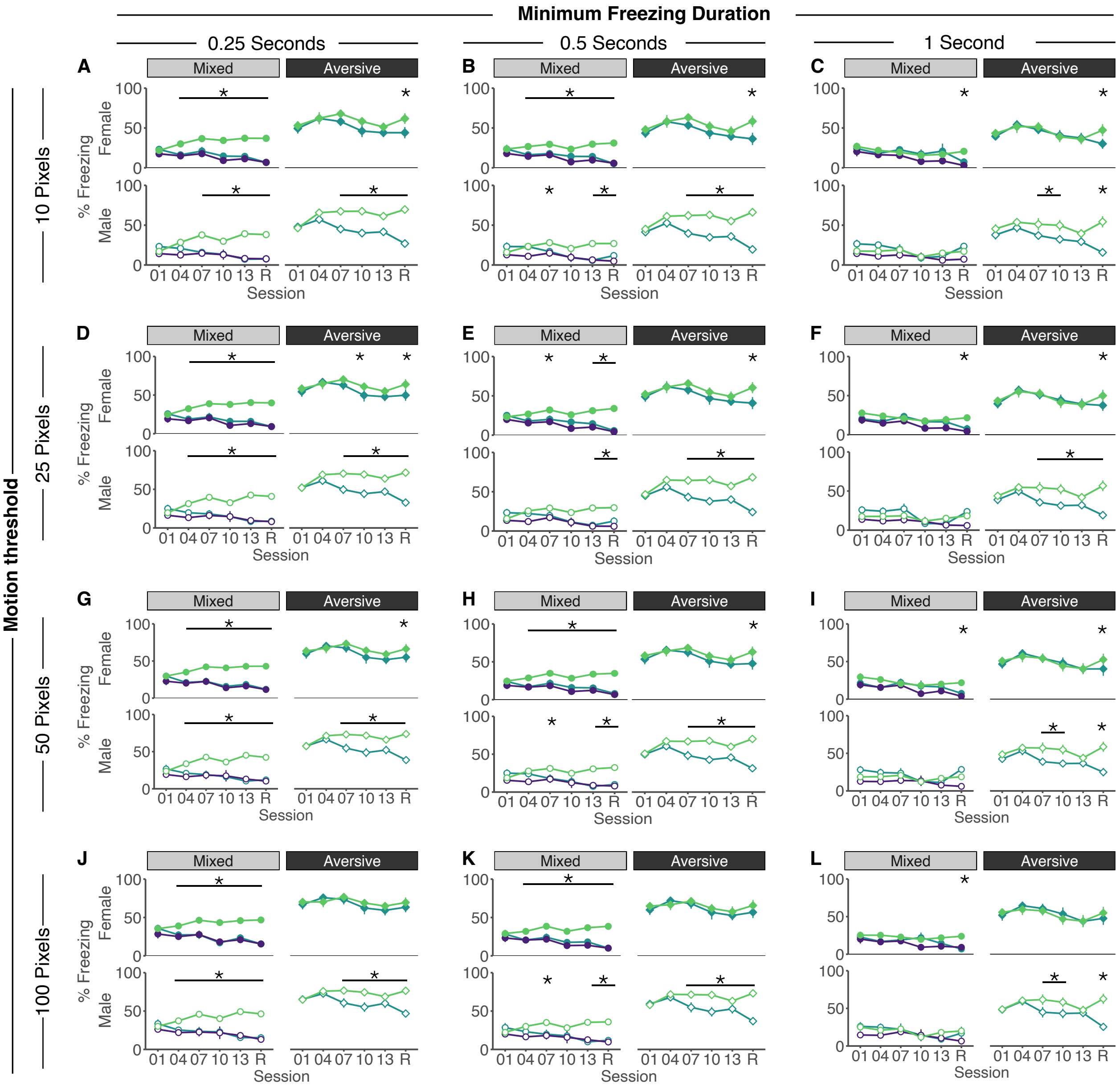


*Figure S3. % Freezing quantified using different motion threshold and freezing duration parameters in ezTrack.* Across all motion thresholds, better discrimination between CS$^-$ and CS$^S$ is observed in the mixed paradigm compared to the aversive only paradigm for females, while males discriminate similarly in both the mixed and aversive only paradigm. With increasing freezing duration, cue discrimination diminishes for both sexes in the mixed paradigm compared to the aversive only paradigm, suggesting shorter freezing bouts in the mixed protocol. * $p_{adj} < 0.05$.

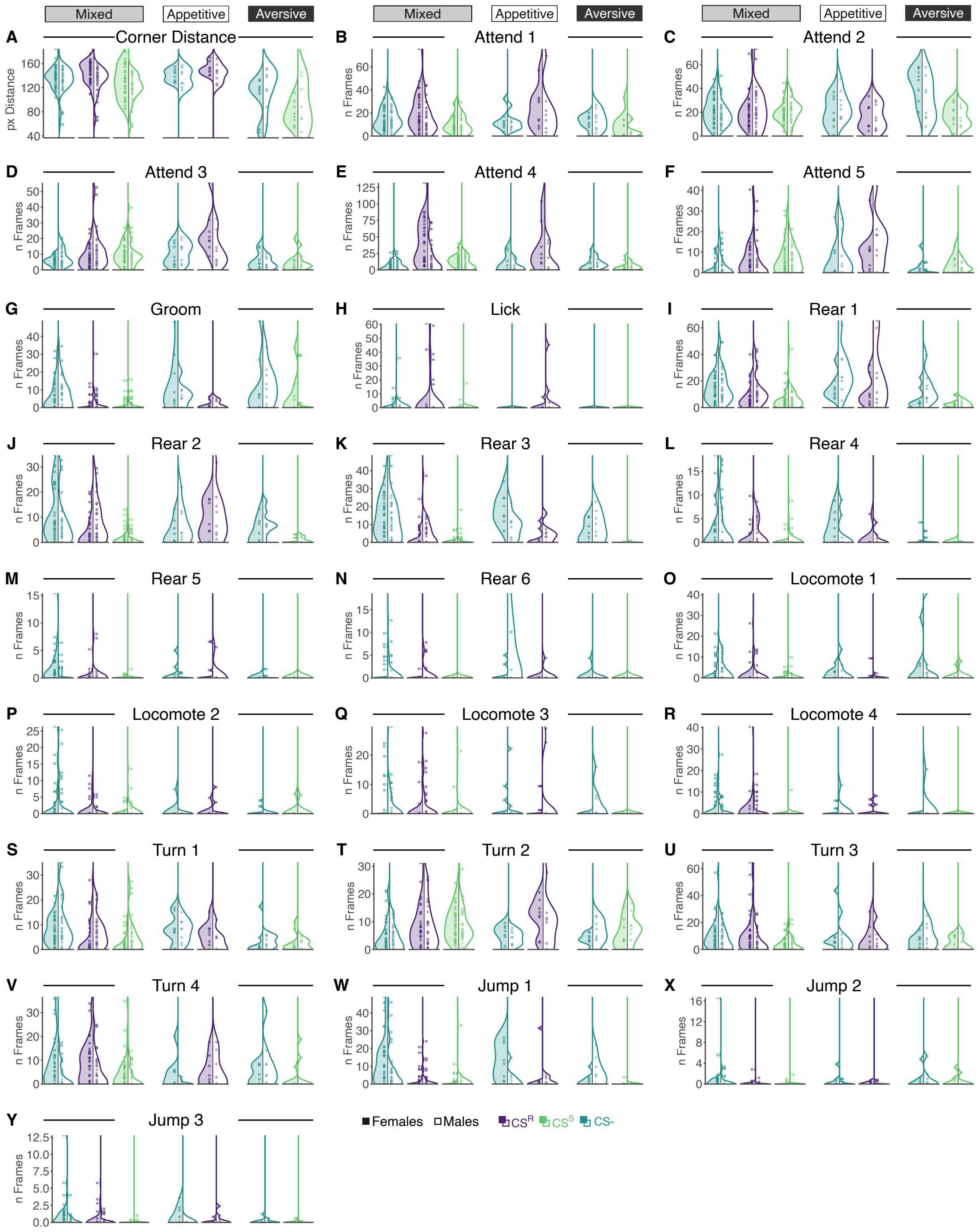


*Figure S4. Raw values on recall day of all variables included in the PLS-DA model not shown in Main Figure 2*. This includes location-based metrics (**A**), attending (**B-F**), grooming (**G**), licking (**H**), rearing (**I-M**), locomoting (**O-R**), turning (**R-V**), and jumping (**W-Y**). Mixed train and test dataset are combined for this visualization.

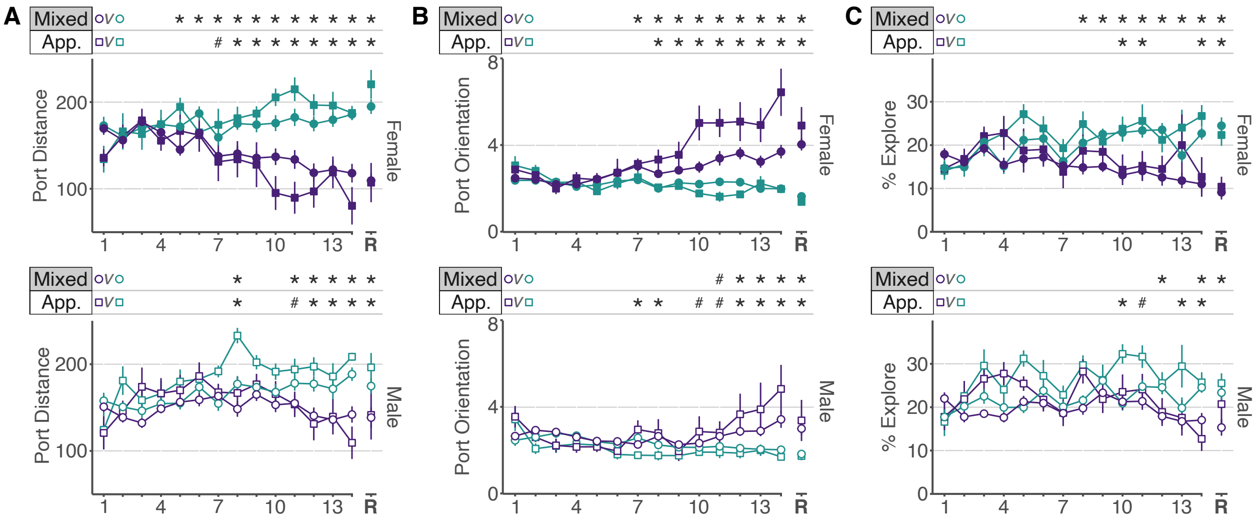


*Figure S5. CS$^R$ predictive behaviors in females (top) and males (bottom)*. (**A**) Mice decrease distance to food port in both mixed and appetitive paradigms, but females do so earlier than males. (**B**) Similarly, port orientation increases in both paradigms, with discrimination being expressed earlier in females than in males. (**C**) % exploration decreases during the CS$^R$ robustly by the end of training in females but not in males. Full statistical results are reported in *Table S8*. * $p_{adj} < 0.05$
